# The potential of *Pseudomonas fluorescens* SBW25 to produce viscosin enhances wheat root colonization and shapes root-associated microbial communities in a plant genotype dependent manner in soil systems

**DOI:** 10.1101/2023.07.21.550058

**Authors:** Ying Guan, Frederik Bak, Rosanna Catherine Hennessy, Courtney Horn Herms, Christine Lorenzen Elberg, Dorte Bodin Dresbøll, Anne Winding, Rumakanta Sapkota, Mette Haubjerg Nicolaisen

## Abstract

Microorganisms interact with plant roots through colonization of the root surface i.e. the rhizoplane or the surrounding soil i.e. the rhizosphere. Beneficial rhizosphere bacteria such as *Pseudomonas* spp. can promote plant growth and protect against pathogens by producing a range of bioactive compounds, including specialized metabolites like cyclic lipopeptides (CLPs) known for their biosurfactant and antimicrobial activities. However, the role of CLPs in natural soil systems during bacteria-plant interactions is underexplored. Here, *Pseudomonas fluorescens* SBW25, producing the CLP viscosin, was used to study the impact of viscosin on bacterial root colonization and microbiome assembly in two cultivars of winter wheat (Heerup and Sheriff). We inoculated germinated wheat seeds with SBW25 wild-type or a viscosin-deficient mutant, and grew the plants in agricultural soil. After two weeks, enhanced root colonization of SBW25 wild-type compared to the viscosin-deficient mutant was observed, while no differences were observed between wheat cultivars. In contrast, the impact on root-associated microbial community structure was plant genotype specific, and SBW25 wild-type specifically reduced the relative abundance of an unclassified oomycete and *Phytophthora* in Sheriff and Heerup, respectively. This study provides new insights into the natural role of viscosin and specifically highlights the importance of viscosin in wheat root colonization under natural soil conditions and in shaping the root microbial communities associated with different wheat cultivars. Further, it pinpoints the significance of microbial microdiversity, plant genotype and microbe-microbe interactions when studying colonization of plant roots.

## INTRODUCTION

Microorganisms associated with plant roots can influence plant health positively through multiple mechanisms e.g. nutrient acquisition, pathogen control and induction of plant defense responses (1). While promising results on exploiting plant growth promoting bacteria to replace or reduce the use of fertilizers and pesticides in agriculture have been obtained (2–4), the translational power from laboratory studies to field performance is currently low (5). This is partly due to the high complexity of natural systems and partly due to our inadequate understanding of processes *e.g.* plant-microbe and microbe-microbe interactions and chemical communication, involved in the bacterial colonization of plant roots. Furthermore, root colonization has primarily been explored in sterile root systems that lack indigenous soil microorganisms (6–8). This ignores the three-way interaction among the inoculant, the root and the indigenous soil community, and competition for colonization space operating under natural soil conditions (9–11). To enhance the success rate of translation from laboratory to field, disentangling genes and processes involved in root colonization in soil systems is of pivotal importance.

Plants and microbes have co-evolved in complex settings for millions of years giving rise to intimate plant-microbe and microbe-microbe interactions, influenced by both soil type, plant age and plant genotype (12–14). Some of these specific interactions involve specialized metabolites (also known as specialized metabolites or natural products) used for both chemical warfare as well as mediators of specific interactions (15, 16). Cyclic lipopeptides (CLPs) produced by *Pseudomonas* sp. are specialized metabolites reported to be involved in root colonization by influencing traits like motility and biofilm formation (17–19), and may thereby also play a role in shaping root-associated microbial communities due to competition for space. Specifically, the CLP viscosin enhances spreading of the producing strain on sterile roots and is essential for motility through swarming (20). In a previous study, we found that viscosin-producing *Pseudomonas* strains are enriched in the rhizoplane of winter wheat in a plant-cultivar dependent manner (Herms et al., unpublished). This suggests a role of viscosin in plant root colonization and plant-microbe interactions dependent on plant genotype. Viscosin has also been identified as a key molecule in microbe-microbe interactions, as it has demonstrated antimicrobial activity against protozoans and protects *P*. *fluorescens* SBW25 from protozoan predation by *Naegleria americana* in lab-based assays, in addition to superior persistence in soil compared to viscosin-deficient mutants (21). Moreover, viscosin has anti-oomycete properties against the economically important plant pathogens *Phytophthora infestans* (22), and *Pythium* (20), both classified as protists. Hence, viscosin seems to have implications for both microbiome assembly and composition which might have consequential effects on plant growth and health. Hence, understanding the impact of viscosin-producing strains on the root microbial communities of wheat grown in agricultural soil could provide a conceptual model to reach higher translational power from laboratory to field.

To unravel these intimate interactions at the root-soil interface and determine the importance of microbial specialized metabolites in early root colonization and microbiome assembly under complex conditions, we used *P. fluorescens* SBW25 as a viscosin-producing model strain in combination with two winter wheat cultivars, Sheriff and Heerup (Heerup naturally enriching for viscosin producing pseudomonads compared to Sheriff; Herms et al, unpublished). We inoculated wildtype *P. fluorescens* SBW25 (SBW25 WT), as well as its corresponding mutant deficient in viscosin production (Δ*viscA*) on wheat seedlings and evaluated their colonization potential and impact on bacterial and protist community assembly in the rhizoplane. We propose two hypotheses: 1) the ability to produce viscosin increases root colonization in a plant-genotype dependent manner and 2) the viscosin-producing strain impacts the protist community by lowering the relative abundance of potential plant pathogenic oomycetes because of the antiprotist properties of viscosin. In addition, we determine whether the ability to produce viscosin has a plant-genotype dependent impact on the microbial community assembly. Finally, we determined plant phenotypes, i.e. biomass, height and root morphology, as responses to the microbial inoculation.

## MATERIALS AND METHODS

### Strain construction and culture conditions

Bacterial strains and plasmids used in this study are listed in Table 1. *P. fluorescens* SBW25 wild type (SBW25 WT) (23) and the viscosin-deficient mutant *P. fluorescens* SBW25 (Δ*viscA*) (22) were routinely grown in Luria Broth at 28°C, shaking at 180 rpm (1% tryptone, 0.5% yeast extract and 1% NaCl). A supplement of antibiotics was used at final concentrations of 50 μg ml^−1^ kanamycin, 100 μg ml^−1^ ampicillin or 10 μg ml^−1^ gentamicin (Table 1). Both strains were chromosomally tagged with mCherry by introducing the mCherry delivery plasmid pME9407 and the helper plasmid pUX-BF13 by electroporation and selection by 10 μg ml^−1^ gentamicin as previously described (19).

**Table 1.**
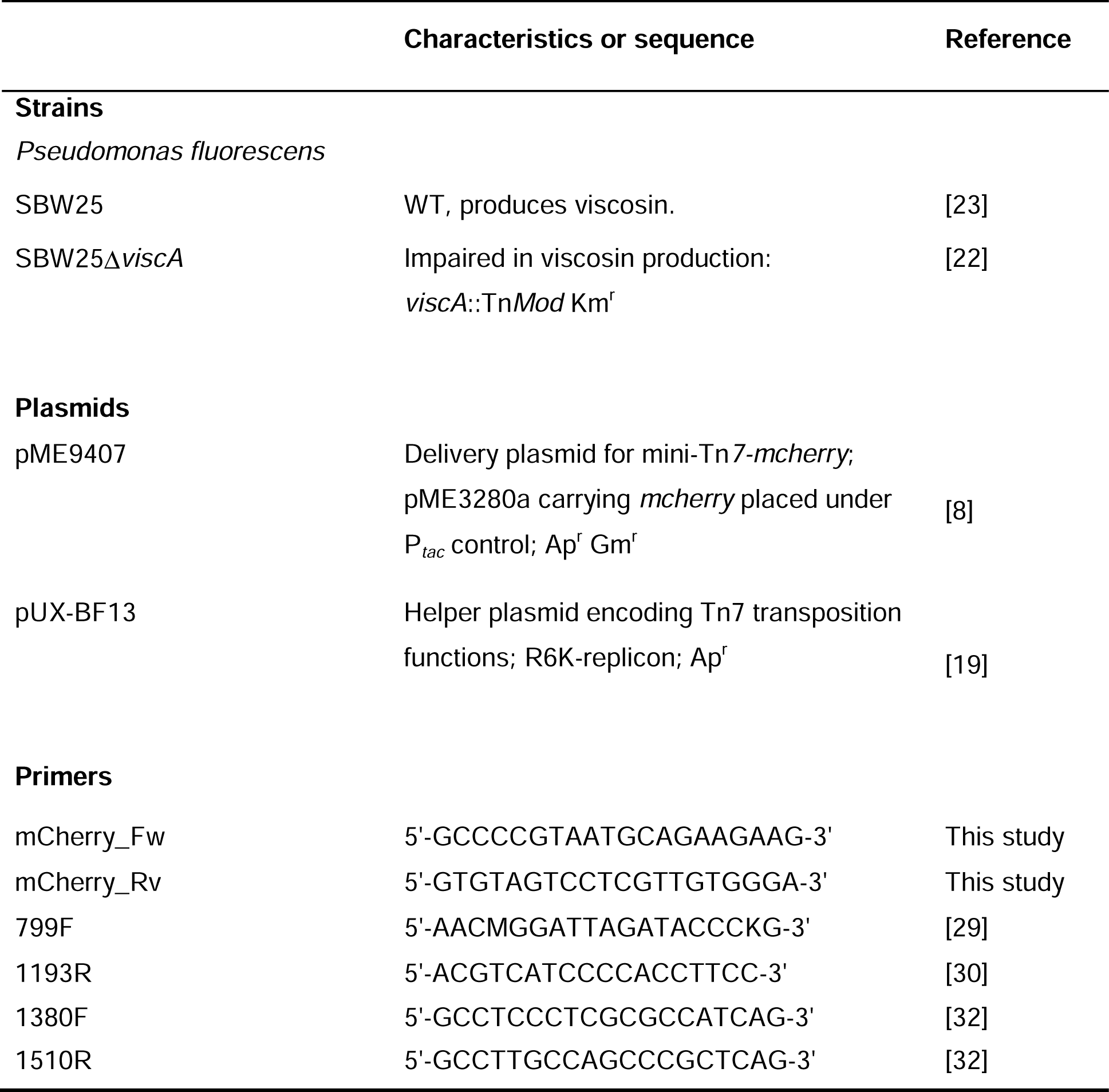
Strains, plasmids and primers.

### Soil collection and properties

Soil was collected from the experimental farm at the University of Copenhagen in Taastrup, Denmark (55° 40′N, 12° 17′E) (24). The soil is a sandy loam (170 g clay kg^−1^, 174 g silt kg^−1^, 362 g fine sand kg^−1^, 255 g coarse sand kg^−1^, and 17 g soil organic matter kg^−1^) (25). Prior to experiments, the soil was air-dried and sieved through a 2 mm mesh.

### Plant experiment setup

Two cultivars of winter wheat, Heerup and Sheriff (Sejet Plant Breeding, Horsens, Denmark), were grown in PVC pots (19 cm high and 3.5 cm in diameter). Soil was thoroughly mixed in the ratio 3:2 with 0.4 - 0.9 mm particle size sand (DANSAND, Brædstrup, Denmark) in a plastic bag. The pots were sealed in the bottom with 50 μm polyamide mesh (Sintab, Oxie, Sweden) before transferring the soil-sand mixture into the pots.

Seeds were soaked in sterile Milli-Q water for 1 h and then transferred to Petri dishes with 2 layers of sterile filter paper, moistened with 5 ml of sterile Milli-Q water. Seeds were kept in the dark at room temperature for 3 days for optimal germination. Overnight *P. fluorescens* cultures were washed twice with sterile 0.9% NaCl, and OD_600nm_ was adjusted to 1.0 (equivalent to approximately 5 × 10^8^ CFU ml^-1^) for seed coating. Germinating seeds with primary roots of 3-4 cm in length were selected and soaked in a Petri dish with bacterial or control solution for 1 hour and immediately transferred to the PVC pots, one seed per pot. The germinated seeds of each wheat species were inoculated as follows: (1) *P. fluorescens* SBW25 mCherry (SBW25 WT); (2) *P. fluorescens* SBW25 Δ*viscA* mCherry (Δ*viscA*) and (3) control treatment with 0.9% NaCl. Eight replicates were set up for each treatment: 5 for DNA extraction and 3 for CFU counting.

The water content of the soil-sand mixture was initially adjusted to 17% (w/w). Each pot was supplemented with 0.83 ml of plant nutrient solution (Drivhusgødning, Park^®^, Schmees, Twistringen, Germany) (NPK 3-1-4), containing 2.04% nitrate-nitrogen (N), 0.3% ammonium-N, 0.46% amide-N, 0.69% phosphorus (P), 4.38% potassium (K), 0.08% sulfur (S), 0.06% magnesium (Mg), 0.033% iron (Fe), 0.013% manganese (Mn), 0.002% copper (Cu), 0.002% zinc (Zn), 0.0006% molybdenum (Mo), and 0.004% boron (B). The plants were grown in a growth chamber under the following conditions: temperature 19/15°C (day/night), relative air humidity 60%/60% (day/night) and photosynthetically active radiation 600/0 μmol m^−2^ s^−1^ (day/night) with a photoperiod of 16h/8h. The pots were watered with deionized water every second day by weighing and watering up to moisture content of 15% throughout the experimental period. Sampling was performed after 14 days of growth. Root sample collection and compartment processing were performed as described by Zervas *et al.* (26) to obtain rhizosphere (soil adhering to the root; exclusively used for confirmation of proper sampling of the compartments as tested by 16S rRNA amplicon sequencing (see below)) and rhizoplane (the root surface) associated microbes, respectively. Samples for DNA extraction were immediately flash frozen in liquid nitrogen and kept on dry ice until storing at −80°C. Samples were freeze-dried (CoolSafe 100-9 Pro freeze dryer, LaboGene, Lynge, Denmark). All freeze-dried samples were stored at −20°C until DNA extraction. Root and shoot length were recorded at sampling, and roots were scanned to determine total root length and root diameter. Following sampling, roots and shoots were dried at 60°C for 3 days before determining the dry weight.

### DNA extraction

Genomic DNA was extracted from 0.5 g sample using a FastPrep-24^TM^ 5G beadbeating system (MP Biomedicals, Irvine, CA, USA) at 6.0 m/s for 40 s and the FastDNA^TM^ SPIN Kit for soil (MP Biomedicals) following the manufacturer’s instructions. The DNA extract was stored at −20°C until further processing for qPCR analysis and rRNA gene amplicon sequencing.

### Quantitative PCR analysis of root colonization ability

Quantification of SBW25 WT and Δ*viscA* was performed by qPCR targeting the mCherry gene using the AriaMx Real-Time PCR (Agilent Technologies, Santa Clara, CA, USA). The primers used are presented in Table 1. Twenty-microliter reactions were prepared with 1×Brilliant III Ultra-Fast SYBR^®^ Green Low ROX qPCR Master Mix (Agilent Technologies, Santa Clara, CA, USA), 1 µg μl^-1^ bovine serum albumin (New England Biolabs^®^ Inc., Ipswich, MA, USA), 0.2 μM of each primer and 2 μl of template DNA. Thermal cycling conditions were as follows: 95°C for 3 min, followed by 40 cycles of 95°C for 20 s and 56°C for 30 s. A dissociation curve was generated at the end of the qPCR program by including a cycle of 95°C for 1 min, 55°C for 30 s, and finally reaching 95°C by increments of 0.5°C s^-1^, each increment followed by a fluorescence acquisition step. Absolute abundance was calculated based on a standard curve for the mCherry target gene (27). Standard curves used for quantification were based on a ten-fold serial dilution of DNA from mCherry-tagged *P*. *fluorescens* SBW25. The standard curve was run with three technical replicates per dilution, and had a dynamic range from 10^2^ to 10^8^ copies/μl. The efficiency ranged from 99.0 to 99.6%, and R^2^ values were > 0.99 for all standard curves.

### CFU analysis of root colonization ability

Colony forming units (CFUs) were determined for rhizoplane samples. Ten-fold dilutions were inoculated on Gould’s S1 (28) agar plates supplemented with 10 μg ml^-1^ gentamicin to select for tagged SBW25 WT and Δ*viscA* cells. Plates were incubated at 28°C in the dark for 48 h before recording CFUs.

### Root imaging and analysis

Prior to root imaging, all roots were thoroughly rinsed to remove soil and sand particles, and stored in distilled water at 4°C. Upon imaging, roots were untangled and arranged in a shallow acrylic dish with distilled water and imaged using an Epson Perfection V700 Photo scanner (Epson, Suwa, Japan) in 8-bit greyscale mode with 400 dpi resolution, subsequently converted to 8-bit JPG for software compatibility. Root length in diameter size classes and average root diameter were determined using the skeletonization method in WinRhizo v. 2016a (Regent Instruments, Quebec, Canada). Roots were divided into 10 size classes between 0 mm diameter and 4.5 mm diameter in 0.5 mm increments and the total root length in each size class was calculated. Total root length, total root surface area, average root diameter and total root volume were also calculated. Size of class fractions were calculated by dividing the root length in a size class by the total root length for each sample pot. Fine roots were determined as those with diameter < 0.5 mm.

Colonization by SBW25 was verified by confocal microscopy targeting the mCherry fluorescent protein. Inoculated seeds were prepared as described above, and then placed in sterile CYG Germination Pouches (Mega International, Roseville, United States). The bags were covered in foil to protect the roots from light. Plants were initially watered with 18 mL sterile water. Three plants per treatment were sat up i.e. for SBW25 and Δ*viscA.* Plants were watered every day with up to 9 mL sterile water to maintain moisture in the bags. Four mL fertilizer was added to each pouch after 1 week. For visualization, seedlings were harvested after one and two weeks, and rinsed in sterile water. For each treatment, three plants were imaged. The roots were sectioned, and for each replicate, a 3-4 cm section was imaged from the top of the root and near the root tip, respectively. Images were obtained by a Leica Stellaris 8 confocal lacer scanning microscope equipped with a supercontinuum white light laser and a HC PL APO CS2 40x/1.25 GLYC objective. Excitation was at 587 nm at a laser power of 2.0%, and emission was collected between 597 nm – 839 nm on a HvD detector with a scan speed of 400 Hz and Line Averages of 6. The gain was between 150-250% to capture mCherry tagged strains. A Trans PMT detector was used to image bright field.

### 16S rRNA and 18S rRNA gene amplicon sequencing

The V5–V7 region of the bacterial 16S rRNA gene was amplified with primers 799F (29) and 1193R (30) (Table 1). This primer pair was found to amplify relatively low proportions of plant mitochondria and chloroplast DNA compared to bacterial DNA (31). The purity and concentration of DNA were determined using a NanoDrop ND-1000 spectrophotometer (Thermo Fisher Scientific, Carlsbad, CA, USA) and a Qubit 2.0 fluorometer (Thermo Fisher Scientific). The ZymoBIOMICS Microbial Community DNA Standard (Zymo Research, Irvine, CA, USA), pure culture SBW25 DNA, and two water controls were included. A two-step dual indexing strategy for Illumina MiSeq (Illumina, San Diego, CA, USA) sequencing was used. First, PCR amplicons were generated in a 25-µl setup using 0.8 U Platinum II Taq (Thermo Fisher Scientific), 1x Platinum II PCR Buffer, 1 mM dNTP mix, 0.2 µM primer 799F, 0.2 µM primer 1193R, and 5 µl DNA template. The PCR thermocycler program included an initial denaturation temperature of 95°C for 2 min, then 33 cycles of 95°C for 15 s, 55°C for 15 s, and 72°C for 15 s and a final elongation step of 72°C for 5 min. The PCR amplification was confirmed by gel electrophoresis on 1.5% agarose gels. PCR products were purified using AMPure XP beads (Beckman Coulter Inc. Brea, CA, USA). The following library construction and Illumina MiSeq sequencing (2 × 300 bp) were performed by Eurofins Genomics (Ebersberg, Germany).

The 1380F and 1510R primer set (Table 1) (32) targeting the V9 region of the 18S rRNA gene was used to evaluate the protist community composition. A two-step dual-indexing strategy for Illumina NextSeq sequencing using a 2-step PCR was used. First, PCR amplicons were generated in 25 µl-reactions with 1x PCRBIO Ultra Mix (PCRBIOSYSTEMS), 0.2 µM of each primer, 0.4 µg μl^-1^ bovine serum albumin (New England Biolabs^®^ Inc., Ipswich, MA, USA) and 5 µl DNA template. The PCR thermocycler program included an initial denaturation temperature of 95°C for 2 min, then 33 cycles of 95°C for 15s, 55°C for 15s, and 72°C for 40 s and finally a final elongation step of 72°C for 4 min. Each reaction in the first PCR was done in duplicates, which were pooled, and used for dual indexing in the second PCR. The second PCR was run with 5 µl of amplicons produced in the first PCR, primers with Illumina adaptors and unique index combinations (i7 and i5) using the reaction conditions described for the first PCR. The second PCR was run using 98°C for 1 min, then 13 cycles of 98°C for 10 s, 55°C for 20 s and 72°C for 40 s and a final elongation of 72°C for 4 min. The PCR amplification was confirmed using 1.5% agarose gels. Subsequently, the amplicon product was cleaned using HighPrep™ magnetic beads (MagBio Genomics Inc. Gaithersburg, MD, USA), according to the manufacturer’s instructions. Finally, DNA concentrations were measured using Qubit 4.0 fluorometer using the High-Sensitivity DNA assay (Thermo-Fischer Scientific). Samples were then equimolarly pooled and sequenced on an Illumina Nextseq sequencer.

### Sequence processing

Raw amplicon reads were processed using the DADA2 pipeline v. 1.14.1 (33). In brief, reads were quality checked and primers were removed using trimLeft in the filterAndTrim function. According to the sequence quality, the 16S rRNA gene reads were filtered using default parameters except for trimRight and minLen (the reads were filtered by truncating the last 20 bp of the forward reads and the last 128 bp of reverse reads using trimRight and minLen = 150 to avoid poor quality and ambiguous sequences). Chimeras were removed after merging denoised pair-end sequences. Each unique amplicon sequence variant (ASV) was assigned to taxa according to SILVA database v. 138.1 (34) and PR^2^ database v. 4.14.0 (35) for the 16S rRNA gene and the 18S rRNA gene, respectively. For bacteria, non-bacterial ASVs including chloroplasts and mitochondrial reads were removed. Similarly, for protist, we discarded plant (Streptophyta), animal (Metazoa), and fungal reads to generate the retained and conservative protist ASV table. To reduce the amount of spurious ASVs, ASVs with a relative abundance below 0.1% in each sample were removed from the dataset (36).

### Data analysis and statistics

Statistical analysis of the plant physiology experiments was performed using GraphPad Prism v. 8.3.0. Differences between two groups were analyzed by unpaired *t*-test (*P* < 0.05). Multiple comparisons were analyzed by one-way analysis of variance (ANOVA) via Tukey’s HSD test (\**P* < 0.05 and \*\**P* < 0.01).

The 16S rRNA and 18S rRNA datasets were analyzed in R version 4.1.3 (37). For microbiome diversity and composition analyses, we used the R packages phyloseq v. 1.38.0 (38), ampvis2 v. 2.7.17 (39) and microeco v. 0.11.0 (40). The amp_rarecurve function in the ampvis2 package was used to generate rarefaction curves (number of reads vs number of observed ASVs) for each sample. For 16S rRNA, we excluded the samples “YG6” (Sheriff), “YG34” (Heerup) and “YG51” (Heerup) from the analysis due to their low read number. Closer inspection of the 18S rRNA samples indicated incomplete removal of rhizosphere soil for sample “YG34” and it was omitted from further analyses. A sample overview is provided in Table S1. The alpha diversity was estimated using Shannon diversity in Divnet v. 0.4.0 (41) with default parameters. This method allows for cooccurrence of taxa in contrast to other methods, which estimate the diversity based on multinomial models. Significance testing of the Shannon diversity was done using beta function in breakaway v. 4.7.6 (42). Samples were not rarefied prior to downstream analyses to avoid discarding information (43).

Principal component analysis (PCA) based on Aitchison distance was performed using the R function ‘prcomp’ and permutational multivariate analysis of variance (PERMANOVA) was used to test the effect of the inoculant treatments, compartment and plant genotype on the beta diversity of the microbial community in vegan v.2.6.2 (44). Analysis of shared and unique ASVs between groups was done using the trans_venn function in microeco package v. 0.11.0 (40).

The differential abundance of ASVs between inoculant treatments (SBW25 WT vs. viscosin deficient mutant-treated, SBW25 WT-treated vs. control, and viscosin deficient mutant-treated vs. control) were analyzed while controlling for the compartment of the two wheat cultivars. The differential abundance was determined using beta-binomial regression with the corncob package v. 0.2.0 (45). Only ASVs that had an estimated differential abundance of □ −1 or >1, and P-values adjusted for multiple testing < 0.05 (FDR < 0.05) were considered significant.

### Data availability

All raw sequencing data used in this study has been deposited in the NCBI Sequence Read Archive (SRA) database under project accession numbers PRJNA928659 (16S rRNA) and PRJNA931264 (18S rRNA).

## RESULTS

### Colonization ability of *P. fluorescens* SBW25 and its viscosin-deficient mutant

To elucidate the importance of viscosin production for the ability of *P. fluorescens* SBW25 to colonize roots of the wheat cultivars Heerup (observed to naturally enrich for viscosin producing bacteria) and Sheriff, we inoculated seedlings with mCherry-tagged *P. fluorescens* SBW25 wild type (SBW25 WT) or its viscosin-deficient mutant (Δ*viscA*). Samples taken immediately after seedling inoculation showed similar colonization potential independent of the ability to produce viscosin, as measured by qPCR and CFU counts, respectively (Fig. 1AB). Furthermore, there was no significant difference in colonization of the two wheat genotypes. After two weeks of seedling growth, qPCR data showed a 4.7-fold higher abundance of SBW25 WT compared to Δ*viscA* in the rhizoplane of Sheriff (*p* = 0.04), whereas a higher, but no significant difference, was observed for the Heerup cultivar (*p* = 0.20) (Fig. 1C). The CFU assay only detected SBW25 WT in the rhizoplane samples from Heerup and Sheriff, whereas no CFUs were observed for Δ*viscA* at the measured dilution, suggesting at least ten-fold higher colonization by SBW25 WT compared to Δ*viscA* (Fig. 1D).

**Fig. 1.**
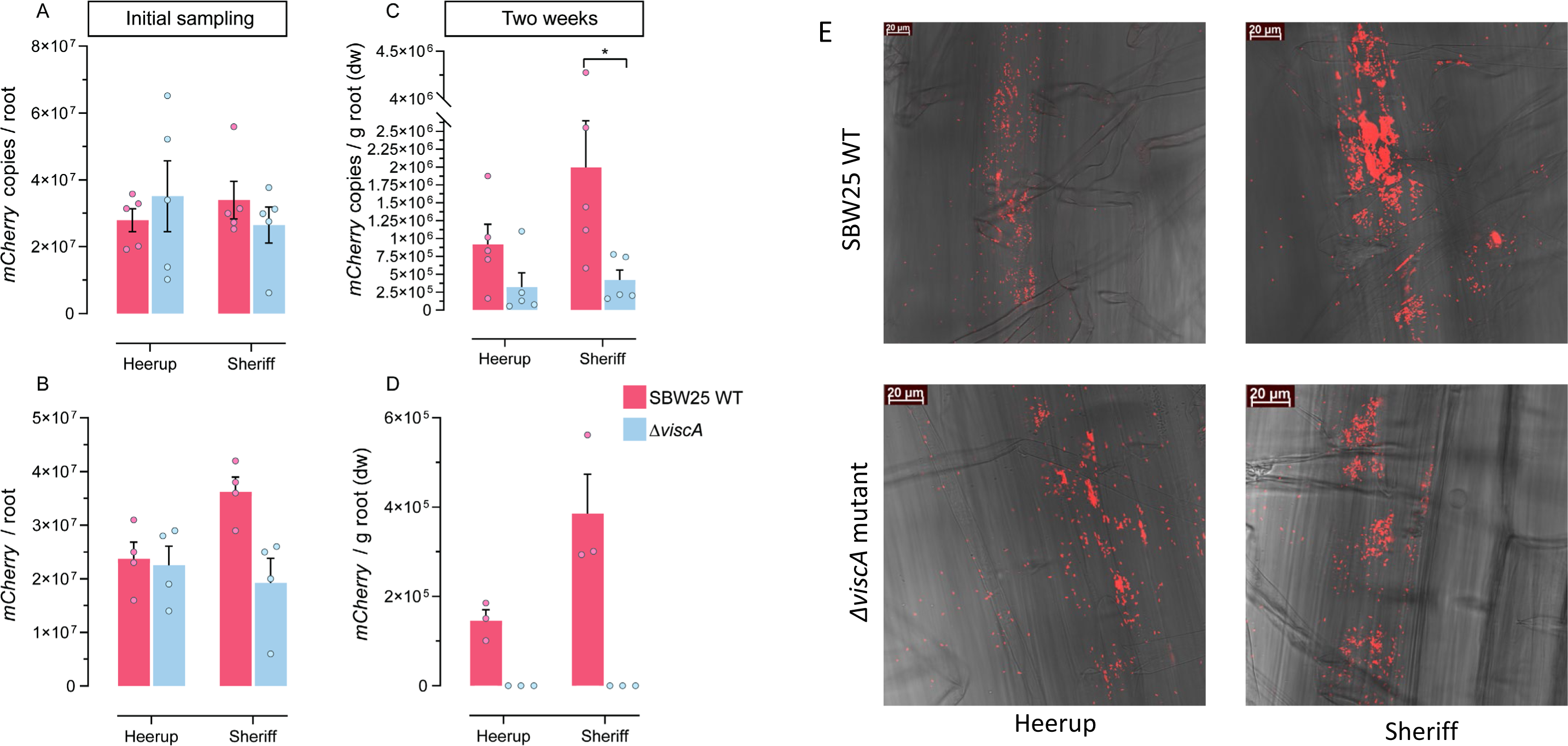
Colonization ability of *P. fluorescens* SBW25 WT and *P. fluorescens* SBW25 Δ*viscA*. (A) qPCR enumeration of inoculated bacteria immediately after inoculation (n=5). (B) CFU enumeration of inoculated bacteria immediately after inoculation (n=4). (C) qPCR enumeration of inoculated bacteria in the rhizoplane at two weeks after inoculation (n=5). (D) CFU enumeration of inoculated bacteria in the rhizoplane at two weeks after inoculation (n=3). Bars represent the mean + standard deviation, and each point represents a sample. Asterisks above histograms indicate whether two groups are significantly different (t-test, *p* ˂ 0.05). (E) microscopy of 2 weeks old plants inoculated with mCherry-tagged *P. fluorescens* SBW25 WT and *P. fluorescens* SBW25 *ΔviscA*, respectively.

In addition to quantitative measures by qPCR, the ability of *P. fluorescens* SBW25 to colonize wheat roots was shown by microscopic detection of cells based on fluorescence emitted by their mCherry-tag (Fig. 1E). Even though the imaging was not quantitative, microscopy supported colonization by both SBW25 WT and Δ*viscA* on the wheat roots.

### Wheat genotype dependent effects of inoculation on plant growth

To evaluate the effect of SBW25 WT and Δ*viscA* on plant growth, we measured shoot and root length as well as shoot and root biomass and performed image analysis of the roots.

Inoculation with SBW25 WT and Δ*viscA* on Sheriff reduced the shoot length with 8% and 6%, respectively, compared to control (*p* < 0.05), whereas no effect was observed for Heerup as compared to the control (Fig. 2A). Root dry weight of Sheriff increased with 53% and 40% after inoculation with SBW25 WT and Δ*viscA*, respectively, compared to the control (*p* < 0.01) Fig. 2B. In opposition, for Heerup, the SBW25 WT and Δ*viscA* had contrasting effects on root dry weight, where Δ*viscA* inoculation increased the root dry weight 27% and 35% compared to the SBW25 WT (*p* □ 0.05) and the control treatment (*p* □ 0.01), respectively (Fig. 2B). We did not observe any effect on root length or shoot dry weight for any of the inoculations as compared to the control treatment (Fig. 2CD). Root image analysis (Fig. S1A) showed a two-fold increase in root tip counts in Heerup (*p* □ 0.05) when inoculated with either SBW25 WT or Δ*viscA* as compared to the control (Fig. S1B, Table S2). Inoculation with SBW25 WT or Δ*viscA* did not affect any of the measured parameters using root imaging for Sheriff. Taken together, the impact of the ability to produce viscosin on plant parameters was dependent on the plant cultivar, as impacts on Sheriff were found to be independent on the ability to produce viscosin, whereas impacts on Heerup were dependent on the ability to produce viscosin. In addition, there was a genotype dependent impact of SBW25, independent of viscosin production, on the development of root tips.

**Fig. 2.**
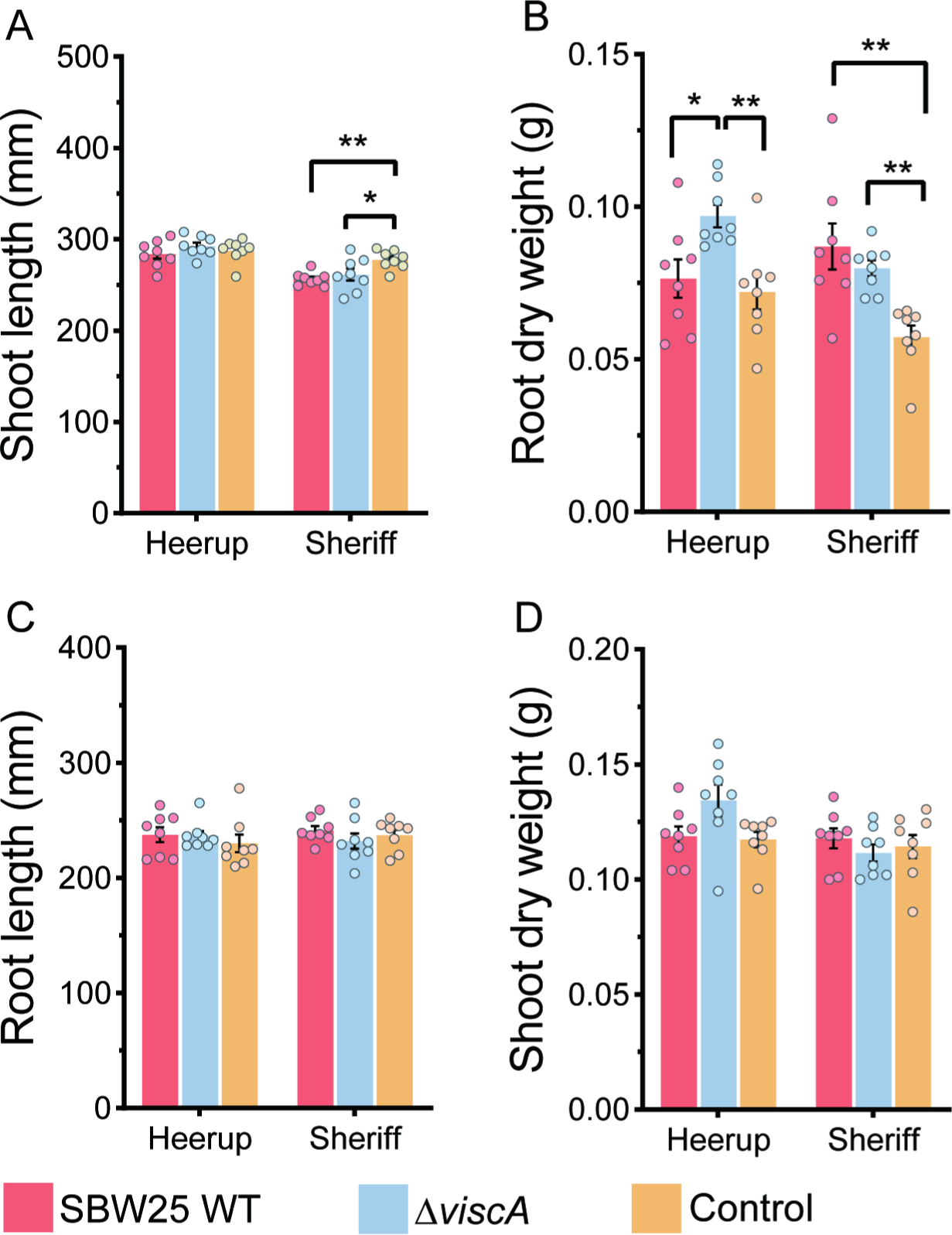
The effect of inoculation with *P. fluorescens* SBW25 WT or *P. fluorescens* SBW25 *ΔviscA* on plant growth (n = 8). (A) Shoot length. (B) Root dry weight. (C) Root length. (D) Shoot dry weight. Bars represent the mean + standard deviation, and each point represents a sample. Asterisks above histograms indicate whether two group are statistically significantly different as assessed by one-way ANOVA followed by a Tukey HSD test: \**P* < 0.05, \*\**P* < 0.01.

### Effects on bacterial community assembly

We used 16S rRNA gene amplicon sequencing, to investigate the effects of SBW25 WT and Δ*viscA* on the rhizoplane bacterial microbiome assembly. The sequencing depths of all samples were sufficient since the number of ASVs was saturated for each sample in the rarefaction curves (Fig. S2A). There were 57 samples with 3 263 907 reads in the total dataset after filtering. The sample sizes ranged from 41 099 to 80 057 reads, with a median of 56 564. The dataset consisted of 449 ASVs. A clear separation of the bacterial communities in the rhizosphere samples from that in the rhizoplane samples (PERMANOVA, *p* < 0.001; R^2^=0.20) confirmed a successful separation of the compartments during sampling (Fig. S3A, Table S3).

The inoculation treatment explained 12% of the variation in the bacterial community composition in the rhizoplane (PERMANOVA, R^2^ = 0.12, *p* = 0.001; Table S4), whereas the interaction between cultivar and inoculation treatment explained 9% of the variation (*R*^2^ = 0.09, *p* □ 0.05; Table S4), indicating a cultivar dependent impact of the inoculation treatments on the community composition. Indeed, ordination visualization shows that there is clear separation of the Δ*viscA* communities from communities resulting from the other two treatments in Heerup (Fig. 3A). In contrast, communities treated with either SBW25 WT or Δ*viscA* clustered apart from the control in Sheriff (Fig. 3B).

**Fig. 3.**
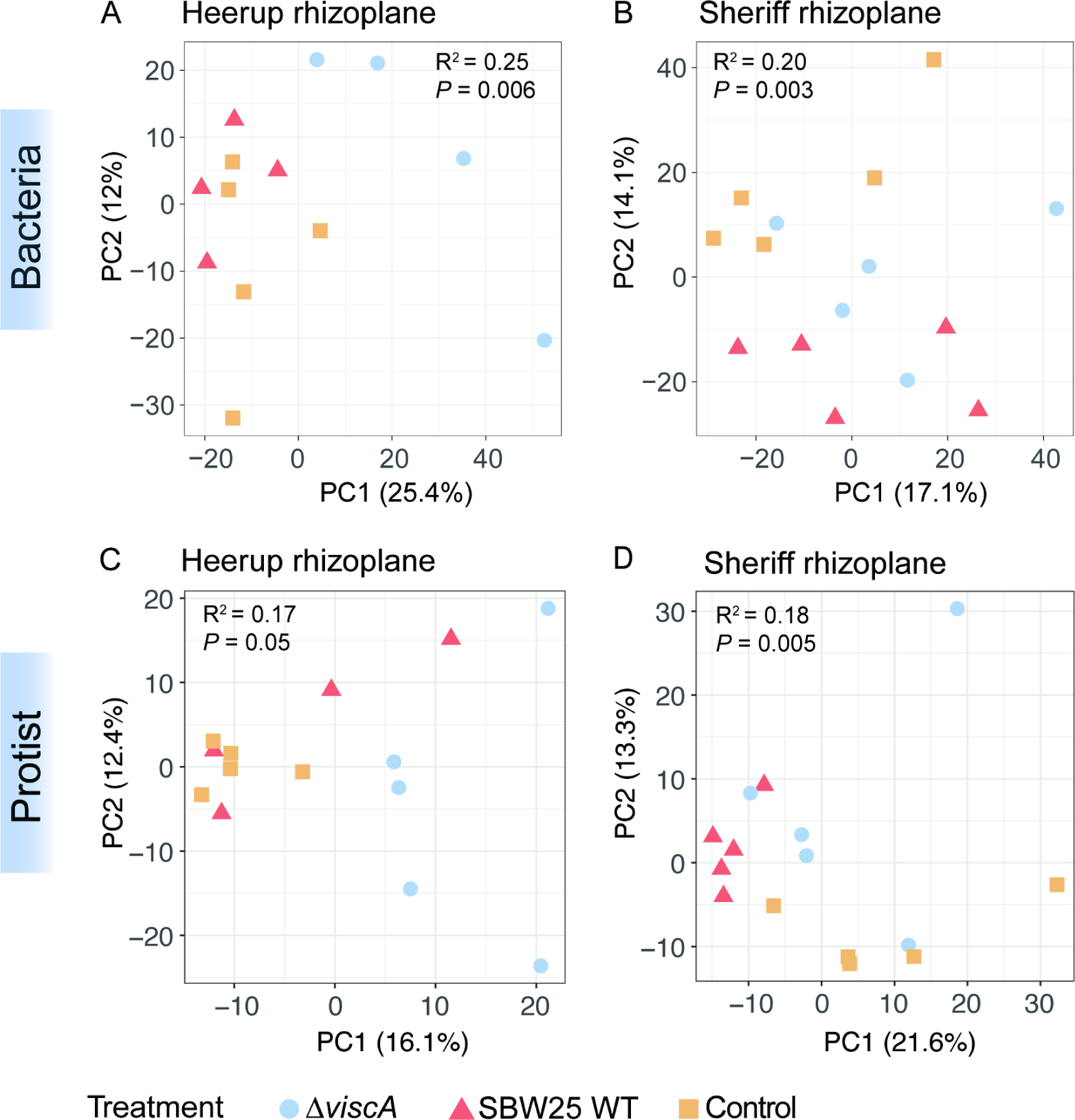
Principal Component Analysis (PCA) based on Aitchison distances of bacterial and protist communities in each subset. (A) Bacteria from the Heerup rhizoplane. (B) Bacteria from the Sheriff rhizoplane. (C) Protist from the Heerup rhizoplane. (D) Protist from the Sheriff rhizoplane. The models were validated using an ANOVA-like permutation test (999 permutations) as indicated by the P-value. R^2^ is expressed as the proportion of the mean sum of squares obtained from PERMANOVA. Each symbol represents individual sample points and samples are colored for inoculation treatment.

Inoculation with Δ*viscA* increased the Shannon diversity (Breakaway, *p* □ 0.001) in the rhizoplane of the Heerup cultivar as compared to the SBW25 WT and the control treatments (Fig. S4A). In the rhizoplane of Sheriff, the Shannon diversity was higher upon inoculation with both the SBW25 WT and Δ*viscA* as compared to the control treatment (*p* □ 0.001). Hence, the influence on bacterial alpha and beta diversity reflects the impact observed on the shoot length and root dry weight (Fig. 2AB), with a similar impact from the SBW25 WT and Δ*viscA* on the Sheriff cultivar, and a differential impact of the two strains on the Heerup cultivar as compared to the control treatment.

### Bacterial community composition response

The rhizoplane communities in both cultivars were dominated by Proteobacteria, Actinobacteriota, and Firmicutes independent of treatment (Fig. S3C). Additionally, *Massilia*, *Paenibacillus*, *Bacillus*, *Dyella*, and *Noviherbaspirillum* were the five most abundant genera across all samples (Fig. S5A).

In the Sheriff rhizoplane, inoculation of SBW25 WT increased (beta-binomial model, *p* □ 0.05) the relative abundance of 14 ASVs and decreased the relative abundance of 11 ASVs compared to the control treatment (Fig. 4A). In contrast, inoculation by Δ*viscA* affected only seven ASVs (*p* □ 0.05), leading to increased relative abundance of three ASVs, and decreased relative abundance of four ASVs compared to the control treatment (Fig. 4B). Comparing the effects of Δ*viscA* inoculation with SBW25 WT inoculation, the relative abundance of five ASVs decreased, whereas the relative abundance of only one ASV significantly increased (*p* < 0.05) (Fig. 4C). Four of the six ASVs with changed relative abundance in the SBW25 treatment compared to Δ*viscA* were impacted in the same way when comparing the SBW25 treatment with the control treatment. These results indicate both a general impact of inoculation with *P. fluorescence* SBW25, and a specific impact based on the ability to produce viscosin in the Sheriff rhizoplane.

**Fig. 4.**
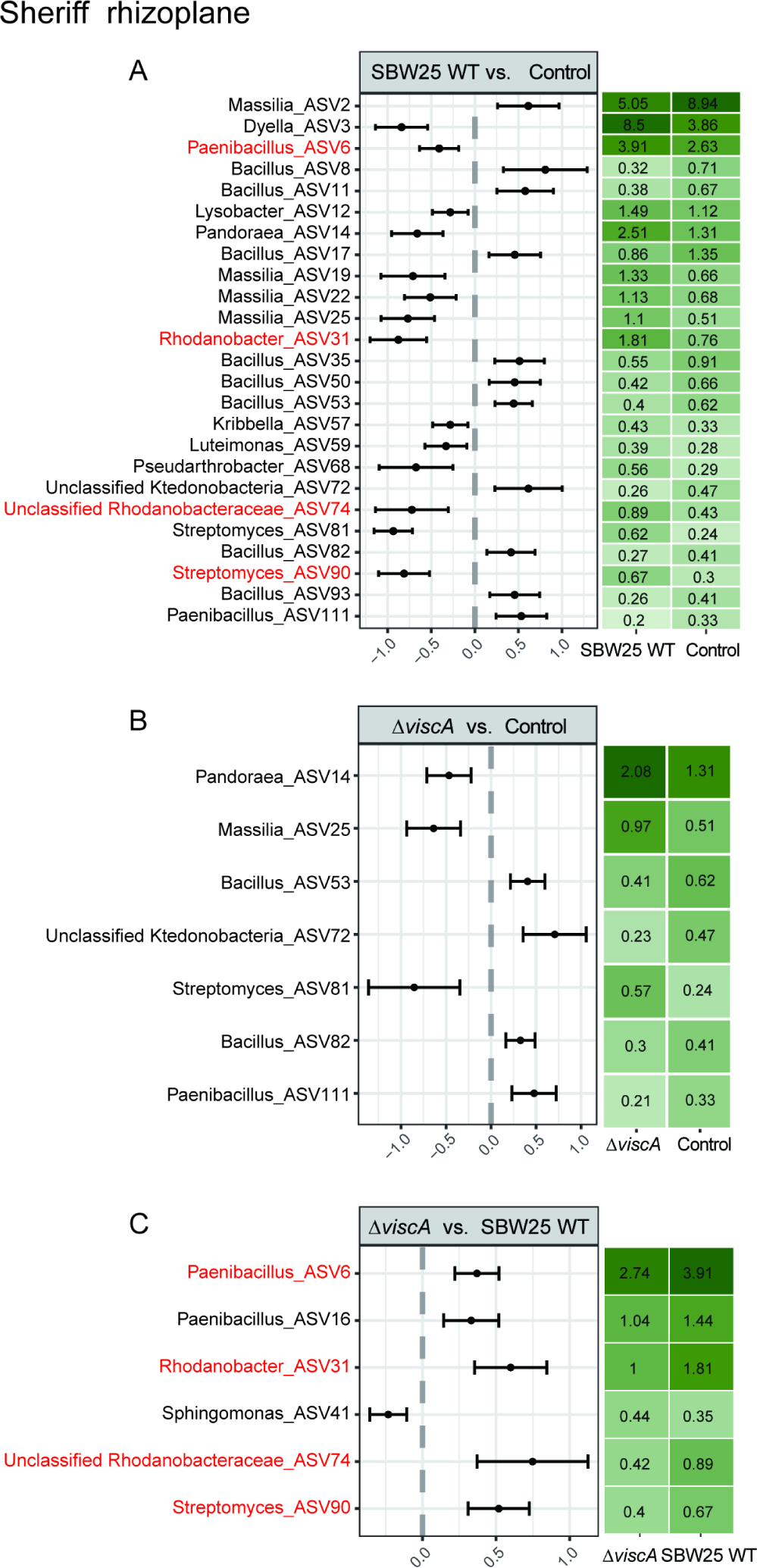
The bacterial ASVs/genera with significant differences in relative abundance between inoculation treatments in Sheriff. (A) Significant differences in ASVs between the *P. fluorescens* SBW25 WT inoculation and Control groups. (B) Significant differences in ASVs between the *P. fluorescens* SBW25 Δ*viscA* inoculation and Control groups. (C) Significant differences in ASVs between the *P. fluorescens* SBW25 Δ*viscA* and *P. fluorescens* SBW25 WT groups. The differential abundance was determined using beta-binomial regression with the corncob. Only ASVs having an estimated differential abundance of ˂ −1 or > 1 and P-values adjusted for multiple testing < 0.05 (FDR < 0.05) were considered significant. The red colored ASVs indicate that inoculation with SBW25 WT showed a significant increase in ASVs compared to both control and mutant treatment.

In the rhizoplane of Heerup, no significant difference in ASV abundance was observed between the SBW25 WT treatment and the control treatment. In contrast, inoculation with Δ*viscA* increased the relative abundance of 39 ASVs and decreased the relative abundance of three ASVs when compared both to the SBW25 WT and the control treatment (Fig. 5AB). An additional four ASVs increased and five ASVs decreased in relative abundance when seedlings were inoculated with Δ*viscA* compared to inoculation with SBW25 WT (Fig. 5A).

**Fig. 5.**
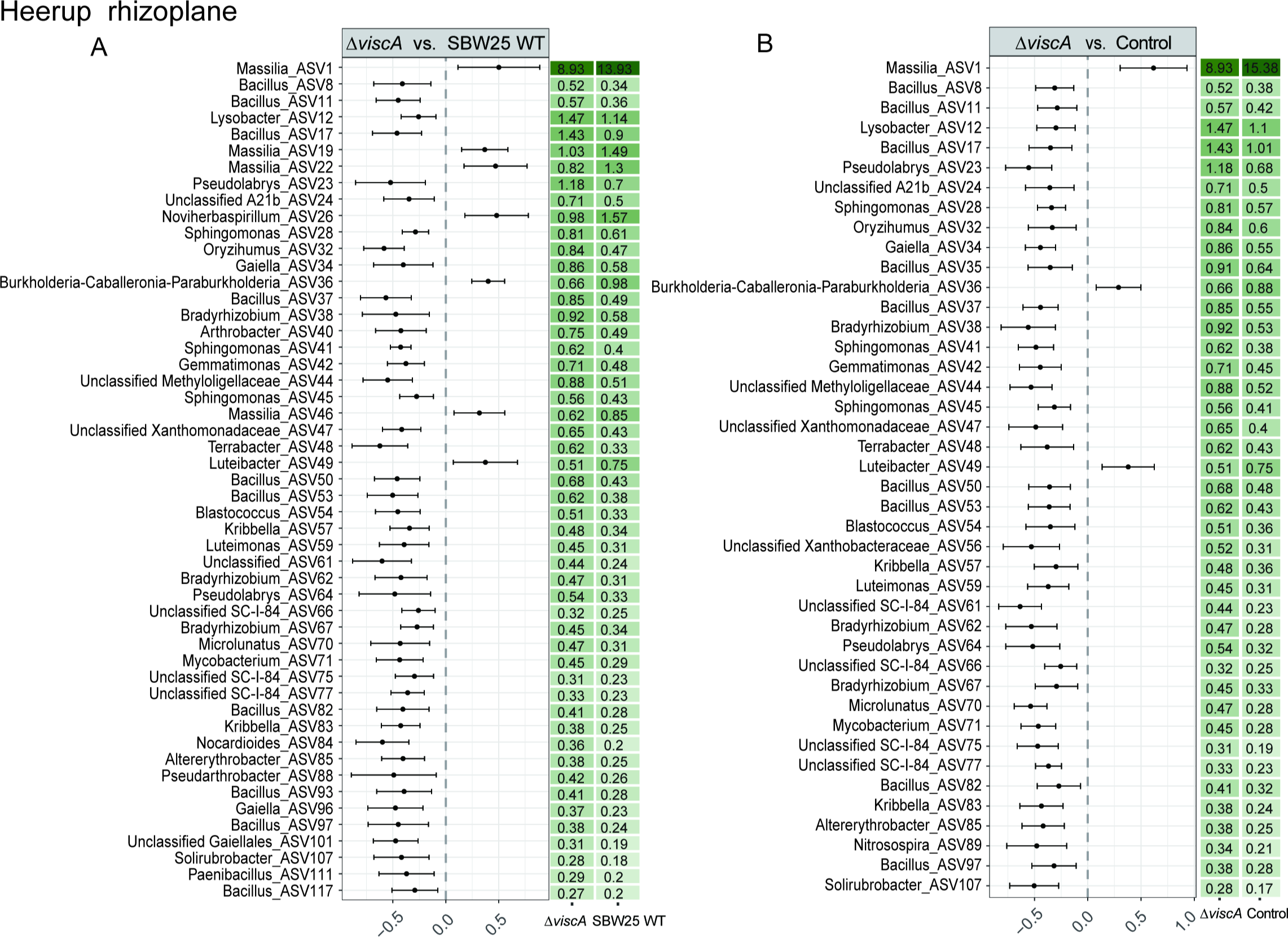
The bacterial ASVs/genera with significant differences in relative abundance between inoculation treatments in Heerup. (A) Significant differences in ASVs between the *P. fluorescens* SBW25 Δ*viscA* and *P. fluorescens* SBW25 WT groups. (B) Significant differences in ASVs between the *P. fluorescens* SBW25 Δ*viscA* inoculation and Control groups. No significant difference in ASVs was found between SBW25 WT treatment and control treatment in Heerup rhizoplane. The differential abundance was determined using beta-binomial regression with the corncob. Only ASV that had an estimated differential abundance of ˂ −1 or > 1 and P-values adjusted for multiple testing < 0.05 (FDR < 0.05) were considered significant.

For both cultivars, ASVs belonging to the genera *Bacillus* and *Massilia* were affected as a response to inoculation treatment. In the Sheriff rhizoplane, inoculations reduced the relative abundance of *Bacillus* and increased the relative abundance of most *Massilia* compared to the control treatment, however more pronounced for SBW25 WT (Fig. 4). In Heerup, Δ*viscA* increased the relative abundance of *Bacillus* and decreased the relative abundance of *Massilia* compared to the SBW25 WT and control treatment. (Fig. 5). Hence, viscosin seems to have a generally negative effect on *Bacillus* ASVs, and a positive effect on the colonization potential of *Massilia* ASVs.

### Effects on protist community assembly

The protist communities were analyzed using 18S rRNA gene amplicon sequencing. The sequencing depth of all samples was sufficient since the number of ASVs was saturated for each sample in the rarefaction curves (Fig. S2B). There were 59 samples with 2 272 445 reads in the total dataset after filtering. The read sizes ranged from 19 495 to 66 875, with a median of 37 300. The dataset consisted of 594 ASVs. Regarding the communities of protists, the compartment was the most important factor for differences in community composition (PERMANOVA, *R^2^* = 0.086, *p* = 0.001) (Fig. S3B, Table S4), again supporting a reliable sampling strategy for obtaining specific compartment samples. Treatment was the second most important factor (R^2^ = 0.04, *p* = 0.009) and thus explained more of the variation in protist community composition than the wheat cultivar (R^2^ = 0.02, *p* = 0.023) (Table S4). As we observed for bacteria, the treatment had a different impact depending on wheat cultivar (Treatment:Cultivar, R^2^ = 0.04, *p* =0.033) (Table S4).

When the cultivars were analyzed individually, the control treatment grouped separately from the SBW25 WT and Δ*viscA* in Sheriff, while the SBW25 WT and the control grouped together in Heerup, resembling the patterns for the bacterial community (Fig. 3CD). For the Sheriff rhizoplane community, the Shannon diversity was lower in SBW25 WT-treated plants than Δ*viscA*-treated plants and control plants (*p* □ 0.001) (Fig. S4B). In contrast, inoculation with SBW25 WT or Δ*viscA* had no effect on the Shannon diversity in the rhizoplane communities of the Heerup cultivar. These results contradict the findings from the bacterial community. One explanation could be the specific impact of SBW25 WT on the oomycete community, with lower impact on other protist community members.

### Protist community composition response

At the division level, the rhizoplane community in both cultivars was dominated by Oomycota (also referred to as Pseudofungi), Cercozoa, and Chlorophyta (Fig. S3D). At the genus level, the five most abundant genera across all samples were *Phytophthora* and four unclassified Oomycota (Fig. S5B). In the rhizoplane of Sheriff, the Cercozoa *Group-Te* was reduced (*p* □ 0.05) following inoculation with SBW25 WT compared to the other two treatments. In addition, inoculation of SBW25 WT reduced (*p* □ 0.05) the relative abundance of an unclassified Oomycota (Fig. 6AB) as compared to the control treatment. No significant difference in ASVs was found between treatment with the Δ*viscA* as compared to the control treatment.

In the rhizoplane of Heerup, one ASV belonging to the Amoebozoa *Leptomyxidae* was increased (*p* □ 0.05) after inoculation with Δ*viscA* compared to the control treatment (Fig. 6C). In contrast, the SBW25 WT treatment reduced the abundance of one ASV belonging to Phytopthora compared to the control, and an ASV from the Rhogostoma lineage when compared with Δ*viscA* treatment (*p* < 0.05, Fig. 6DE). Inoculation with SBW25 WT showed a trend of decreased relative abundance of *Pythium* as compared to Δ*viscA* and control treatments in the Sheriff rhizoplane (Fig. 6F). For the Heerup rhizoplane, the trend was an increase in *Pythium* when inoculated with Δ*viscA* as compared to SBW25 and control treatments, with no difference between the SBW25 and control treatments.

**Fig. 6.**
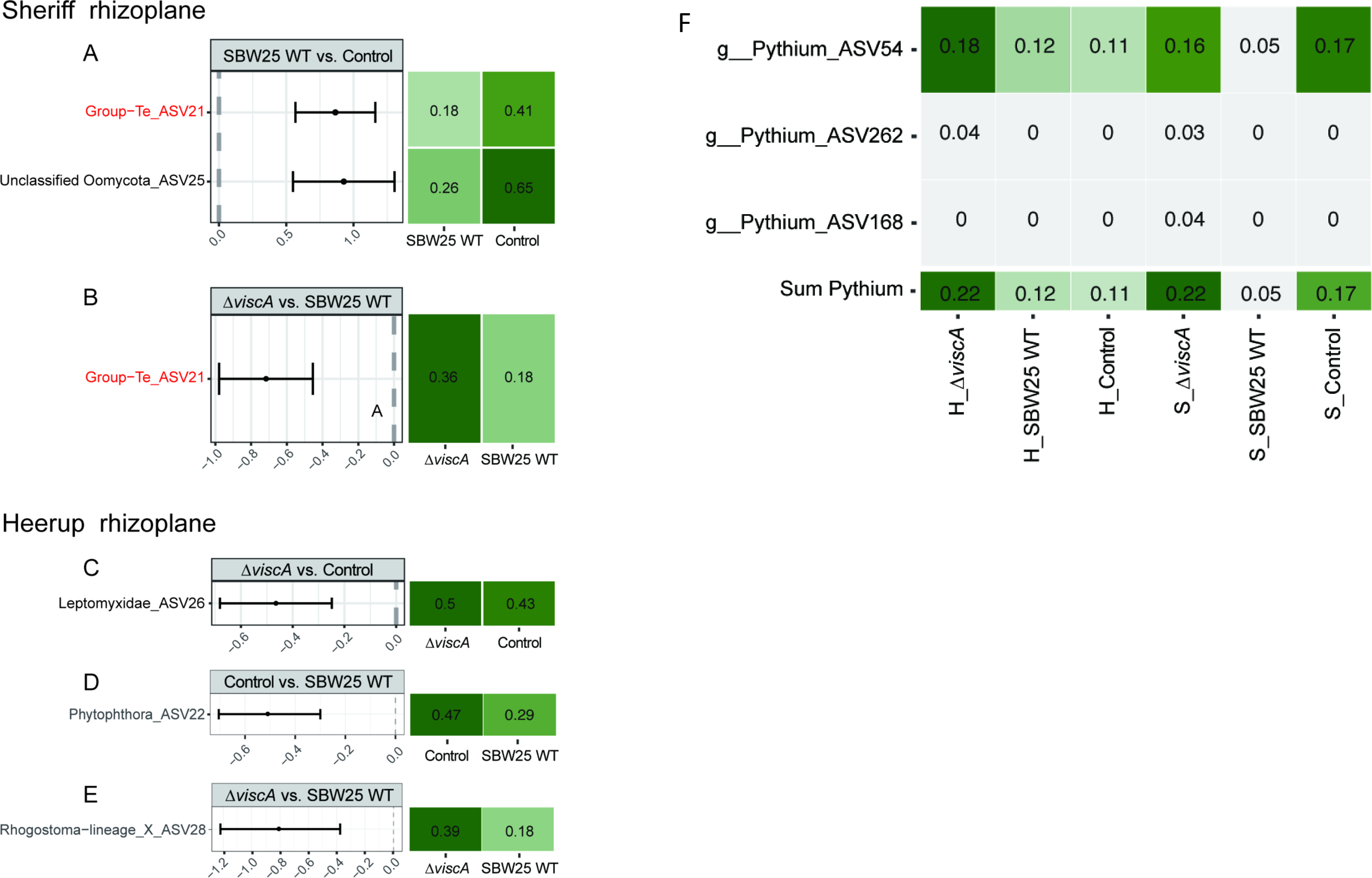
The protist ASVs/genera with significant differences in relative abundance between inoculation treatments in Heerup and Sheriff. (A) Significant differences in ASVs between *P. fluorescens* SBW25 WT inoculated and control groups in Sheriff. (B) Significant differences in ASVs between the *P. fluorescens* SBW25 Δ*viscA* and *P. fluorescens* SBW25 WT groups in Sheriff. (C) Significant differences in ASVs between the *P. fluorescens* SBW25 Δ*viscA* inoculation and Control groups in Heerup. (D) Significant differences in ASVs between the Control and *P. fluorescens* SBW25 WT groups in Heerup. (E) Significant differences in ASVs between the *P. fluorescens* SBW25 Δ*viscA* and *P. fluorescens* SBW25 WT groups in Heerup. In Sheriff rhizoplane samples, no significant differences in ASVs were found between the Δ*viscA* and the control treatment. (F) The relative abundance of *Pythium* in the rhizoplane was analyzed in each group. The mean relative abundance of all *Pythium* in each treatment group (n=5). Sum represents the abundance of these three ASVs for each group. H: Heerup; S: Sheriff. The red colored ASVs indicate that inoculation with SBW25 WT showed a significant decrease in ASVs compared to both control and mutant treatment.

## DISCUSSION

Understanding key drivers of microbial colonization and microbiome assembly at the root-soil interface is fundamental for harnessing the positive effect of beneficial plant-microbe interactions on plant performance. In the present study, we examined the impact of adding a viscosin producing *P. fluorescens* SBW25 WT compared to a viscosin-deficient mutant (Δ*viscA*) to two wheat cultivars: Heerup, observed naturally to enrich for viscosin producing pseudomonads, and Sheriff. Specifically, we studied whether the potential of SBW25 to produce viscosin impacts root colonization and root microbial community assembly in these two cultivars.

The ability to produce viscosin enhanced the colonization potential of SBW25 WT compared with Δ*viscA* in both cultivars, contrasting our first hypothesis and observations that viscosin producing *Pseudomonas* strains are enriched in a culture-dependent manner in the wheat rhizoplane (Herms et al., unpublished). This may be explained by the inoculation strategy applied in our experiment, where inoculants were added by root dipping, giving SBW25 a competitive advantage in colonization over bacteria colonizing from the soil community. Viscosin has previously been shown to be involved in surface spreading on sterilized sugar beet roots in potting compost as well as colonization of broccoli florets (46) and our results expand this finding by demonstrating the importance of viscosin in wheat root colonization in soil during competition with other microorganisms. Furthermore, other CLPs, such as massetolide A and amphisin are important for colonization of tomato roots and sugar beet seeds, respectively (17, 18) pointing towards a natural role of CLPs in root colonization. For viscosin this phenotype of enhanced colonization could be due to the amphiphilic properties of the compound, which alter the surface charge of the bacteria or the root surface for improved colonization (47). In contrast to these findings, Yang *et al.* (48) did not observe any impact on rrhizosphere colonization potential in wheat based on the potential to produce viscosin when comparing *P. fluorescens* HC1-07 and its viscosin-deficient mutant. However, the *P. fluorescens* HC1-07 strain was mutated in the *viscB* gene (48), whereas the impairment of viscosin production in SBW25 used in this study, was due to a mutation in the *viscA* gene (19). Whether these contrasting results are mutant generation, soil condition, plant cultivar or chemically-dependent remains to be elucidated.

The abundance of SBW25 WT and Δ*viscA* on plant roots decreased 100-fold over the course of the experiment and they comprised less than 1% of the total community after two weeks. This is in the range of the *Pseudomonas* genus in rhizosphere samples at a relative abundance of 0.32% across eight winter wheat cultivars and eight soil types from Europe and Africa (49). Other work also demonstrated that despite wheat seed inoculating with growth-promoting *Bacillus* strains they only comprise 2-3% of the seedling community after 4 weeks in the soil (50). This supports the notion that the assembly of the rhizoplane community is highly deterministic after the initial root tip colonization which is more random (51). This leaves only a small part available for exchange of bacteria. Alternatively, the origin of the strains, sugar beet leaf for SBW25 (23), or adaptation to laboratory conditions, could impede their growth in a natural system. Combined, this might account for the often-low transitional power observed when going from simple testing in greenhouse systems not using soil to field performance of bacterial inoculants.

Despite the colonization potential of SBW25 WT on both wheat cultivars, a pronounced difference in the microbial community assembly, measured by ASVs significantly changing between treatments, was observed between the two cultivars. The multitude of ASVs changing in relative abundance, regardless of whether SBW25 WT or Δ*viscA* was introduced in the Sheriff rhizoplane, hint to a general effect of *P*. *fluorescens* SBW25 independent of viscosin production on Sheriff microbiome assembly. Additionally, a specific effect of SBW25 WT was observed, with four ASVs from the genera *Paenibacillus, Rhodanobacter, Streptomyces* and unclassified Rhodanobacteriaceae increasing in relative abundance after the addition of SBW25 WT in comparison to inoculation with Δ*viscA*. These ASVs can be hypothesized to benefit from the viscosin produced by SBW25 WT, either through increased colonization or reduced predators. However, to fully determine the impact of viscosin on specific microbes, viscosin production must be detected in the rhizoplane habitat, which currently is not technically possible at relevant concentrations in soil systems. In contrast to the *P. fluorescens* SBW25 and presumed viscosin effects on the bacterial community assembly identified in the Sheriff cultivar, no ASVs changed significantly in the Heerup rhizoplane as a response to SBW25 WT inoculation. One explanation for the lack of impact seen for the SBW25 WT could be that the Heerup rhizoplane is naturally colonized by viscosin-producing pseudomonads, and that the presence of these strains shape the overall microbial assembly at the roots of this cultivar. Hence, it can be speculated that the viscosin-producing pseudomonads are first colonizers of the Heerup rhizoplane under natural conditions. Indeed, *Pseudomonas* has previously been found to be the most dominant taxa in the active microbial community in 12-days old wheat rhizoplane, suggesting a dominant role in early root community assembly (52). Hence, independent on whether the pseudomonads are soil-derived or inoculated viscosin producers, they play a defined role in the further assembly of the Heerup rhizoplane microbiome. Taken together, viscosin production appears to be important in root community assembly and suggests a role of specialized metabolite production for root community assembly in general. These observations thus highlight the need for future research to evaluate the importance of strain specific specialized metabolites for microbiome assembly in general, and whether such intimate interactions would be dependent on root exudate composition or root architecture and morphology (53). On the other hand, inoculation with the Δ*viscA* mutant revealed a differential impact on the microbial community as compared to the control and the SBW25 WT treatment in Heerup. Interestingly, the majority of affected ASVs increased in relative abundance in the Δ*viscA* mutant treatment, hinting to a competitive advantage of harboring the viscosin gene or an antagonistic effect of viscosin on these taxa.

*Bacillus* and *Massilia*, genera previously shown to be associated with the wheat rhizosphere (49, 54), were observed to be specifically impacted by the inoculations in a wheat cultivar and inoculation dependent manner, collectively accounting for 50% and 25% of the impacted ASVs in Sheriff and Heerup, respectively. *Bacillus* decreased significantly as a response to *P. fluorescens* independent of viscosin in the Sheriff rhizoplane, but with a more pronounced effect by SBW25 WT. *Bacillus* was found to have a significantly higher relative abundance in the rhizoplane of Heerup when Δ*viscA* was inoculated, as compared to both the control and the SBW25 WT treatment. This suggests a direct impact of SBW25 WT on the *Bacillus* community. Since *Bacillus* is well known for its plant-growth promoting abilities (55) the microbe-microbe interactions suggested by the data presented here could have impacts on whether stable establishment of plant-growth-promoting rhizobacteria is successful under natural conditions.

In contrast to *Bacillus*, *Massilia* increased in relative abundance by introduction of SBW25 WT and to a lesser extent Δ*viscA*. Strains belonging to the genus *Massilia* are known as copiotrophic root colonizers (49, 56, 57), and have been proposed as a key member of the wheat root microbiome (49). Furthermore, *Massilia* species are known to colonize the endophytic compartment of wheat, and hence may play an important role for early stage development and microbial assembly (54). In summary, the present study emphasizes the importance of studying specialized metabolites and microbe-microbe interactions in soil systems to gain a full understanding of microbiome assembly at the root-soil interface.

Protists play an important role in the rhizosphere because of their effect on nutrient availability in the soil (58) and impact on the structure of the microbial communities (59), e.g. through predation. Yet, protist communities at the root zones are understudied compared to bacterial and fungal communities (60–62). In the Sheriff and Heerup cultivars, SBW25 WT decreased the relative abundance of a single oomycete ASV (ASV 25) and Phytophthora ASV (ASV 22), respectively. This finding supports our second hypothesis that viscosin-producing bacteria reduce the abundance of oomycetes. Additionally, a decrease in *Pythium* was observed in Sheriff, supporting the above findings. While no single *Pythium* ASV changed significantly between the treatments in Heerup, a trend of increased *Pythium* abundance when plants were inoculated with Δ*viscA* as compared to SBW25 WT and the control treatment was observed. If Δ*viscA* replaces naturally occurring viscosin producers, thereby resulting in fewer *Pythium* antagonists, this could explain the increase in *Pythium* abundance. Oomycetes such as *Phytopthora* and *Pythium* are abundant in the plant-soil habitat (63), and while many species are saprophytes, there are also several plant pathogenic species (64). Hence, viscosin production by microbial inhabitants of the rhizoplane might be important for decreasing abundance of these potential pathogens. In general, the inoculation with SBW25 had only a minor effect on the protist communities, with few ASVs being significantly affected. In the Sheriff cultivar, we observed a decrease of the Cercozoa *Group-Te* in the SBW25 WT treated rhizoplane. *Group-Te* is found in the rhizosphere of multiple crops and model plants, e.g. maize, Arabidopsis, potato (65, 66), but the ecology of this organism is unknown. While the overall findings support a resilience of the microbial communities in the Heerup rhizoplane, the diversity estimates showed a differential pattern for the bacterial and the protist community, respectively, with no difference in the protist diversity measure upon inoculation with either strain of SBW25. This could be explained by the natural recruitment of viscosin producers by Heerup, leading to a minor effect on the protist community despite a possible replacement of the soil-borne viscosin producers by Δ*viscA*. In contrast, a lack of natural recruitment of viscosin producers by Sheriff would explain the lower diversity of protists in the Sheriff rhizoplane community compared to the Δ*viscA* and control treatments in Sheriff. In summary, these findings support a plant genotype specific impact of the ability to produce viscosin on the protist community.

The resilience in the bacterial community of the Heerup rhizoplane was in agreement with the phenotypic response of the plant. Hence, the root dry weight significantly increased when the plant was challenged with the mutant, which also caused significant changes in the microbiome compared to the SBW25 WT and water treatment. For the Sheriff cultivar, both the SBW25 WT and Δ*viscA* caused a significant shift in the microbiome, which coincided with a significant increase in root dry weight for both treatments compared to the control. Whether there is a direct link between viscosin production and root dry weight is currently not known, as it could also be caused by secondary effects from a changing microbiome.

Despite the similar colonization potential of the two wheat cultivars observed for SBW25 WT and Δ*viscA*, respectively, SBW25 WT was found to also impact plant root architecture parameters dependent on plant genotype. It has previously been shown that inoculation of plants with plant beneficial bacteria alters root morphology (67–69), but to our knowledge this is the first time that it has been shown to be cultivar-dependent. The observed differential effect on plant root parameters could be the result of differential community assembly, dependent both on plant genotype and/or the ability of the inoculant to produce viscosin.

In conclusion, the ability to produce viscosin enhances root colonization in both cultivars, contrasting our hypothesis of cultivar-dependent root colonization. Conversely, our second hypothesis was supported as root colonization by SBW25 WT reduced the abundance of potential plant pathogenic oomycetes, including *Phytophtora*, in a cultivar dependent manner. In addition, the relative abundance of multiple bacterial taxa was affected by SBW25 WT colonization in a cultivar dependent manner. Even though factors like soil properties and community composition are important for microbiome assembly in the rhizoplane, this work demonstrates the impact of a specific specialized metabolite on microbial community assembly in the rhizoplane in a plant genotype dependent manner. Acknowledging these plant genotype specific differences is important, and we urge future studies to include several cultivars when investigating root colonization by single strains. This knowledge is important to provide advance our fundamental understanding of microbial ecology in the plant-soil interface and such knowledge can be applied in the future to develop more robust microbial inoculants for plant growth promotion.

## Acknowledgments

Thanks to Dorette Müller-Stöver and Marie Louise Bornø for supplying experimental soils. Thanks to Alex Gobbi for helping prepare the bacterial sequencing library and Athanasis Zervas for helping with the NextSeq. Thanks to our group members Kitzia Yashvelt Molina Zamudio, Jonathan Sølve, and Dorthe Thybo Ganzhorn for their support with the sampling. Imaging data were collected at the Center for Advanced Bioimaging (CAB), University of Copenhagen, Denmark. This study was funded by the Novo Nordisk Foundation (Grant number: NNF19SA0059360), and the Chinese Scholarship Council for a Ph.D. scholarship (CSC Grant 201908510124).

## Competing Interest Statement

The authors declare no competing interests.

## Notes

### Competing Interest Statement

The authors have declared no competing interest.

### Summary of Updates

This version has been revised to focus the story. Additional data has been added in the form of microscopy picture of root colonization.

