## Supplementary Figures for "The potential of *Pseudomonas fluorescens* SBW25 to produce viscosin enhances wheat root colonization and shapes root-associated microbial communities in a plant genotype dependent manner in soil systems"

Fig. S1. Root image analysis. (A) Images of representative samples of the roots. (B) Results of seven parameters (total root length, surface area, average diameter, root volume, tips, fine root length, and thick root length) of the root (n=4). Bars shown in (B) represent mean ± standard deviation, and each point represents a sample. Asterisks indicate statistically significant difference (**P* < 0.05, one-way ANOVA followed by Tukey’s HSD test).

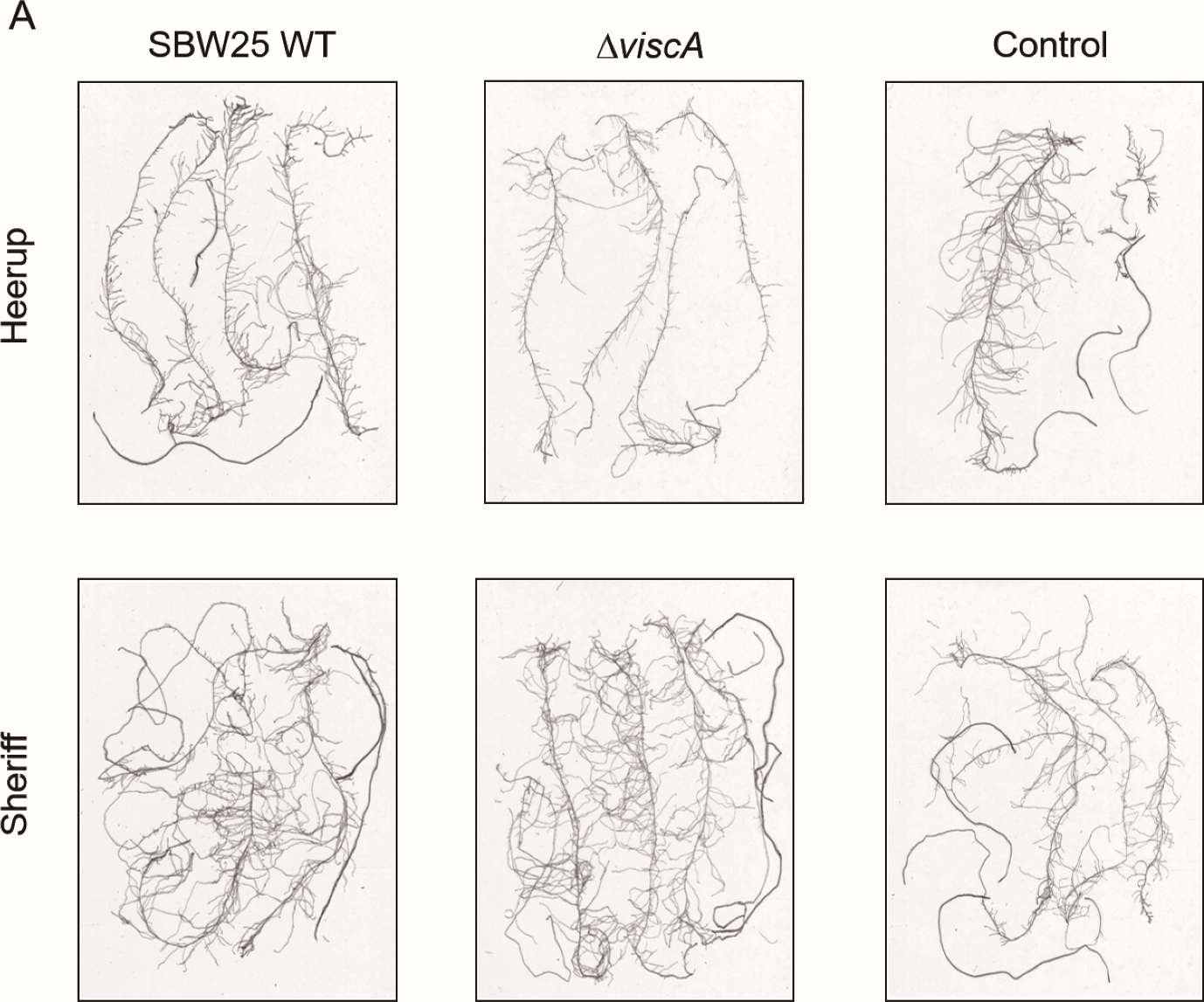

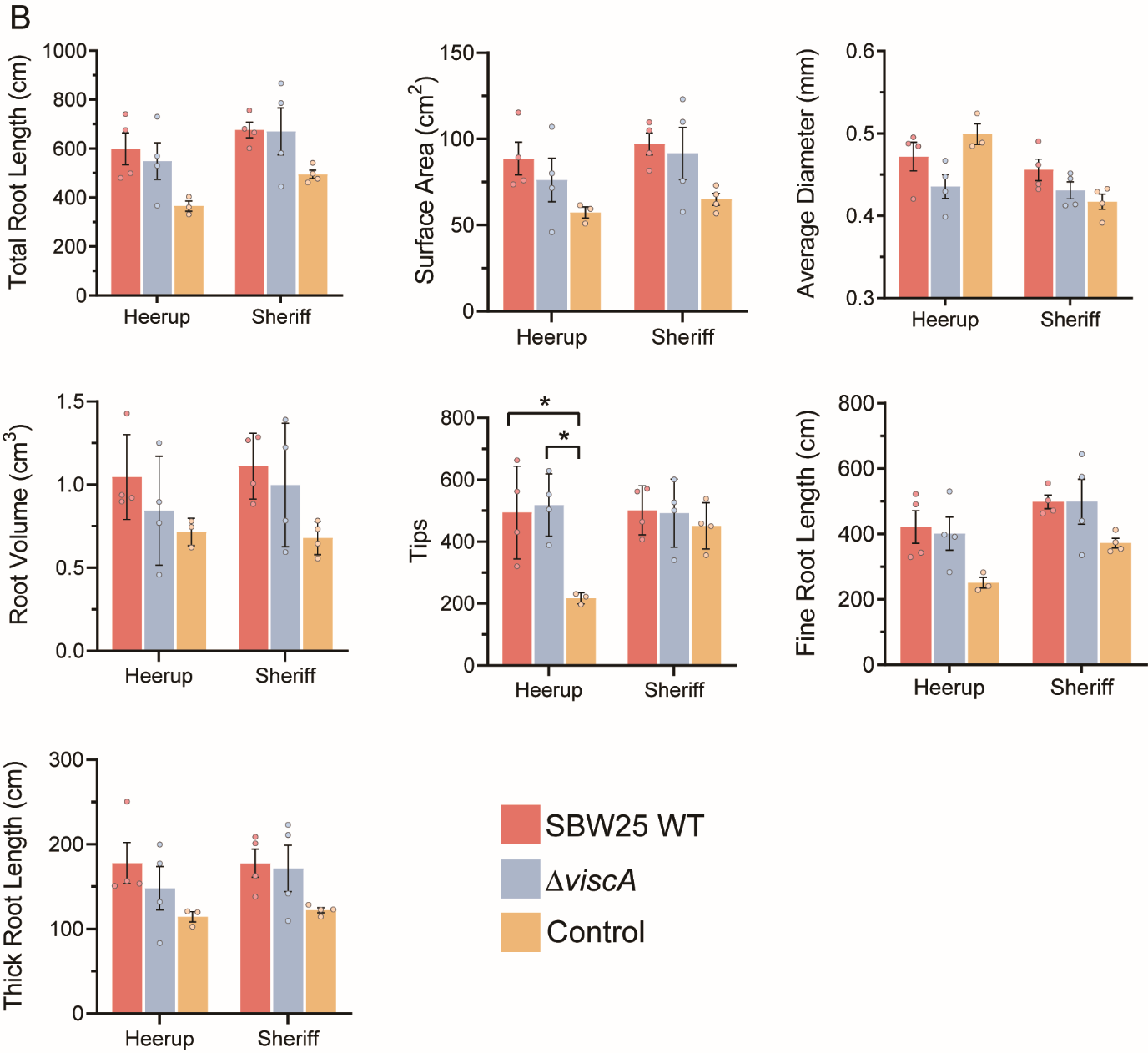

Fig. S2. Rarefaction curves of the 16S rRNA (A) and 18S rRNA (B) gene amplicon reads. Colors represent different compartments (RP: rhizoplane, RS: rhizosphere), cultivars (H: Heerup, S: Sheriff) and inoculation treatments (*viscA*, SBW25 WT and Control).

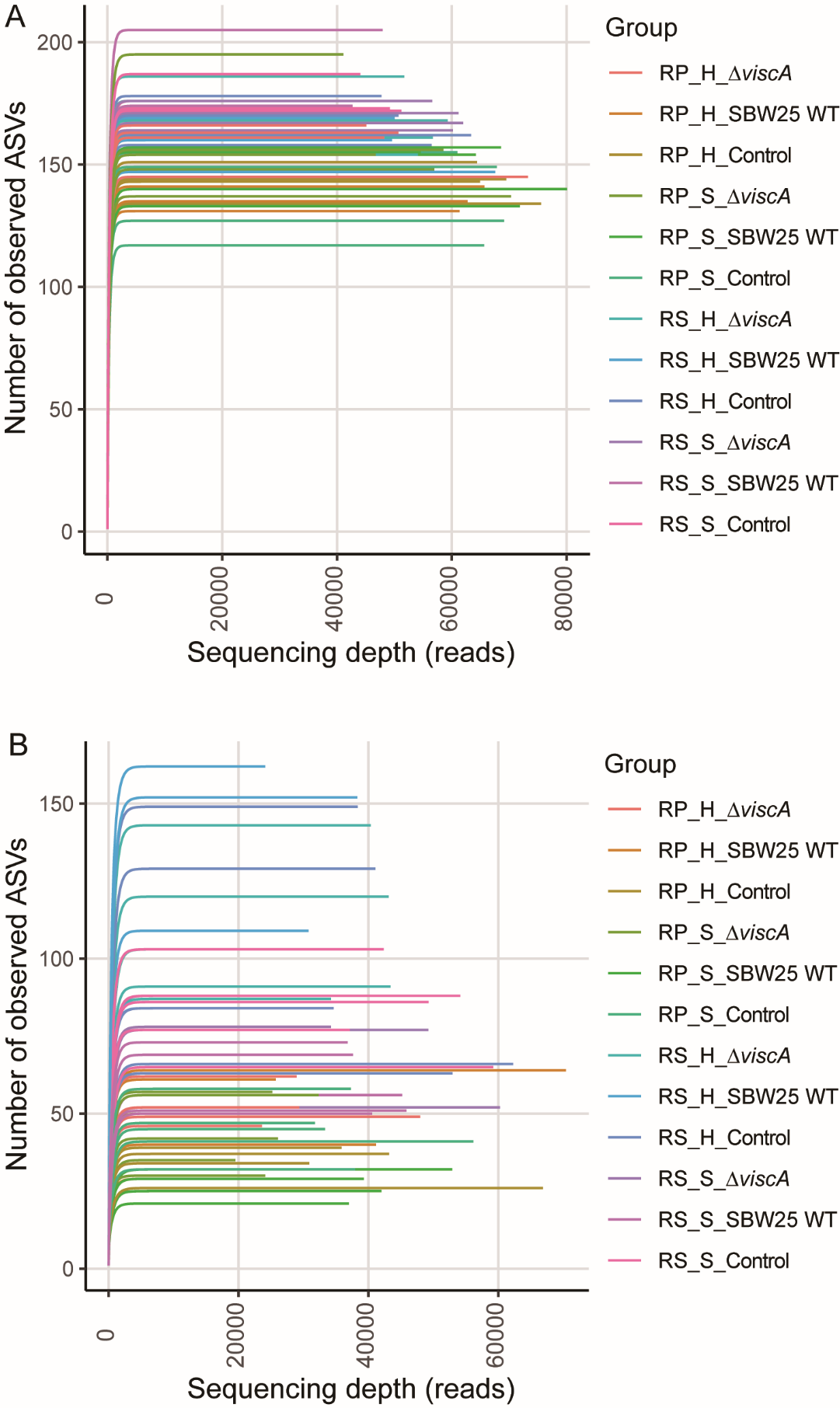

Fig. S3. Principal Component Analysis (PCA) based on Aitchison distances of bacterial and protist communities for all samples. (A) Bacterial community. (B) Protist community. Constrained by compartment. The models were validated using an ANOVA-like permutation test (999 permutations) as indicated by the P-value. R^2^ is expressed as the proportion of the mean sum of squares obtained from PERMANOVA. Each symbol represents individual sample points. (C) Bacterial composition at phylum level; (D) Protist composition at division level. RP: rhizoplane, RS: rhizosphere; H: Heerup, S: Sheriff; SB: SBW25 WT, M: Δ*viscA*, C: Control.

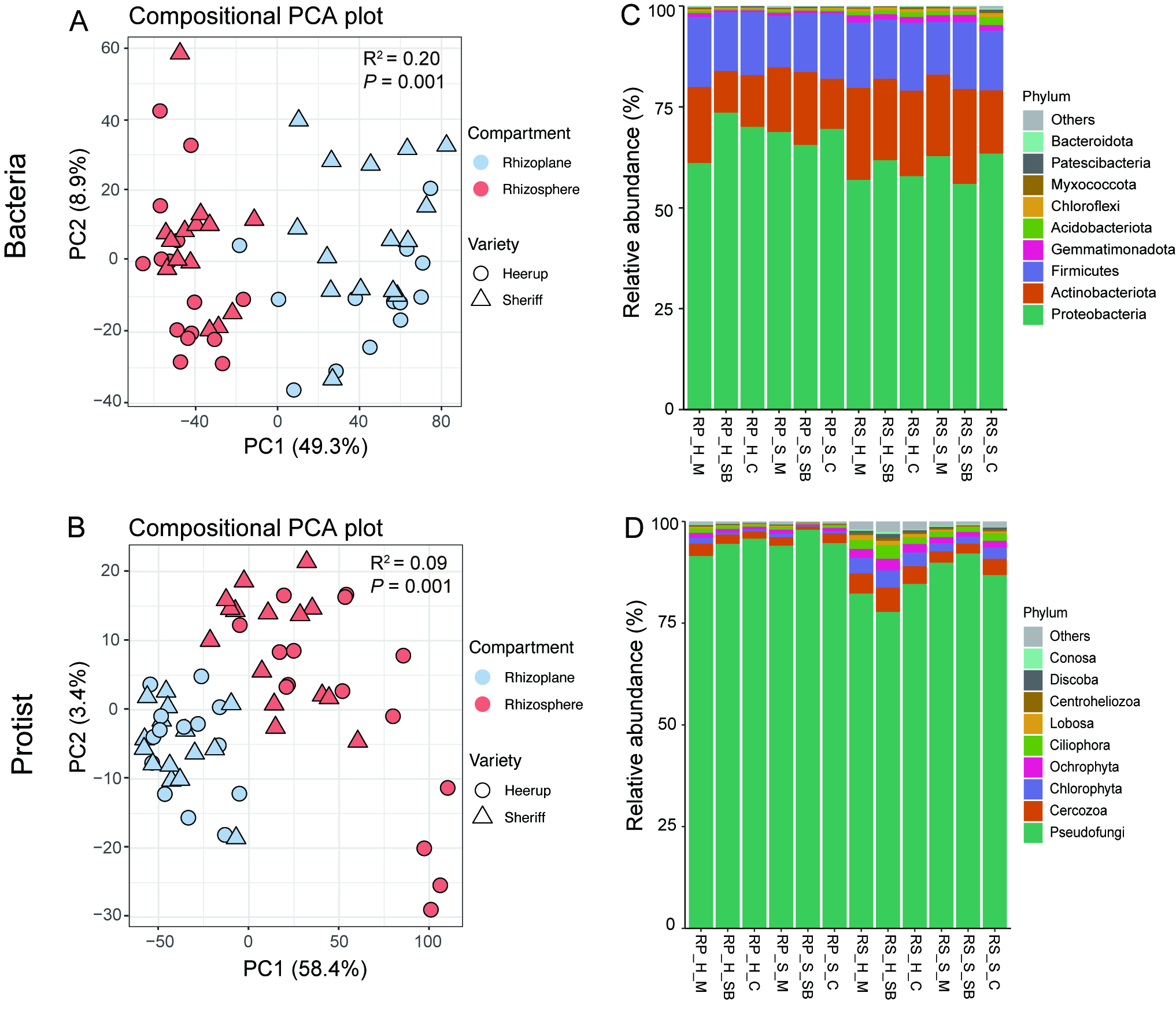

Fig. S4. Shannon diversity estimated at the ASV level. (A) Bacteria; (B) Protist. The mean values and error bars are estimated using Divnet. The significance testing of Shannon diversity was done using ‘betta’ function in breakaway. Different letters indicate significant differences between samples (*P* ˂ 0.05).

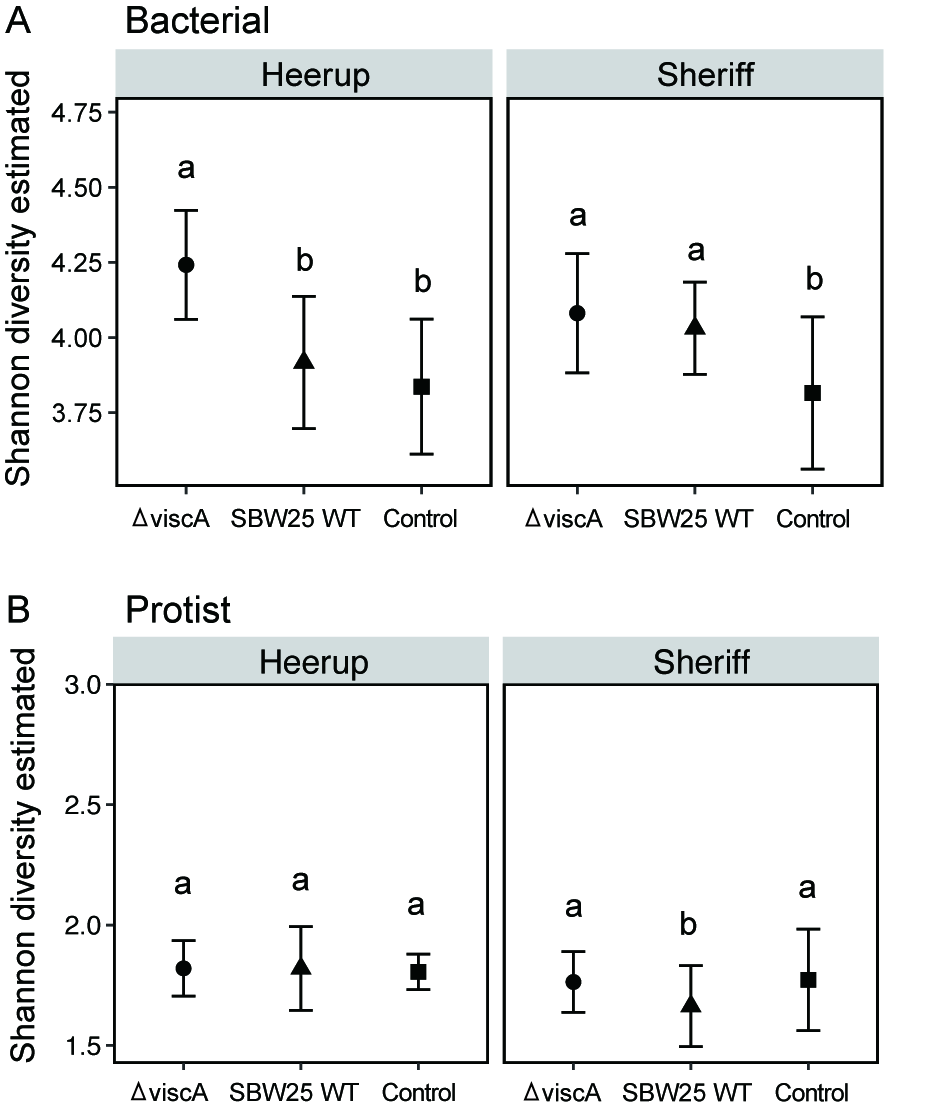

Fig. S5. Heatmap of the 15 most abundant genera across all samples (Include the phylum and class name). Values are average relative abundance (%), n = 5. (A) Bacteria; (B) Protist. RP: rhizoplane; RS: rhizosphere; H: Heerup; S: Sheriff.

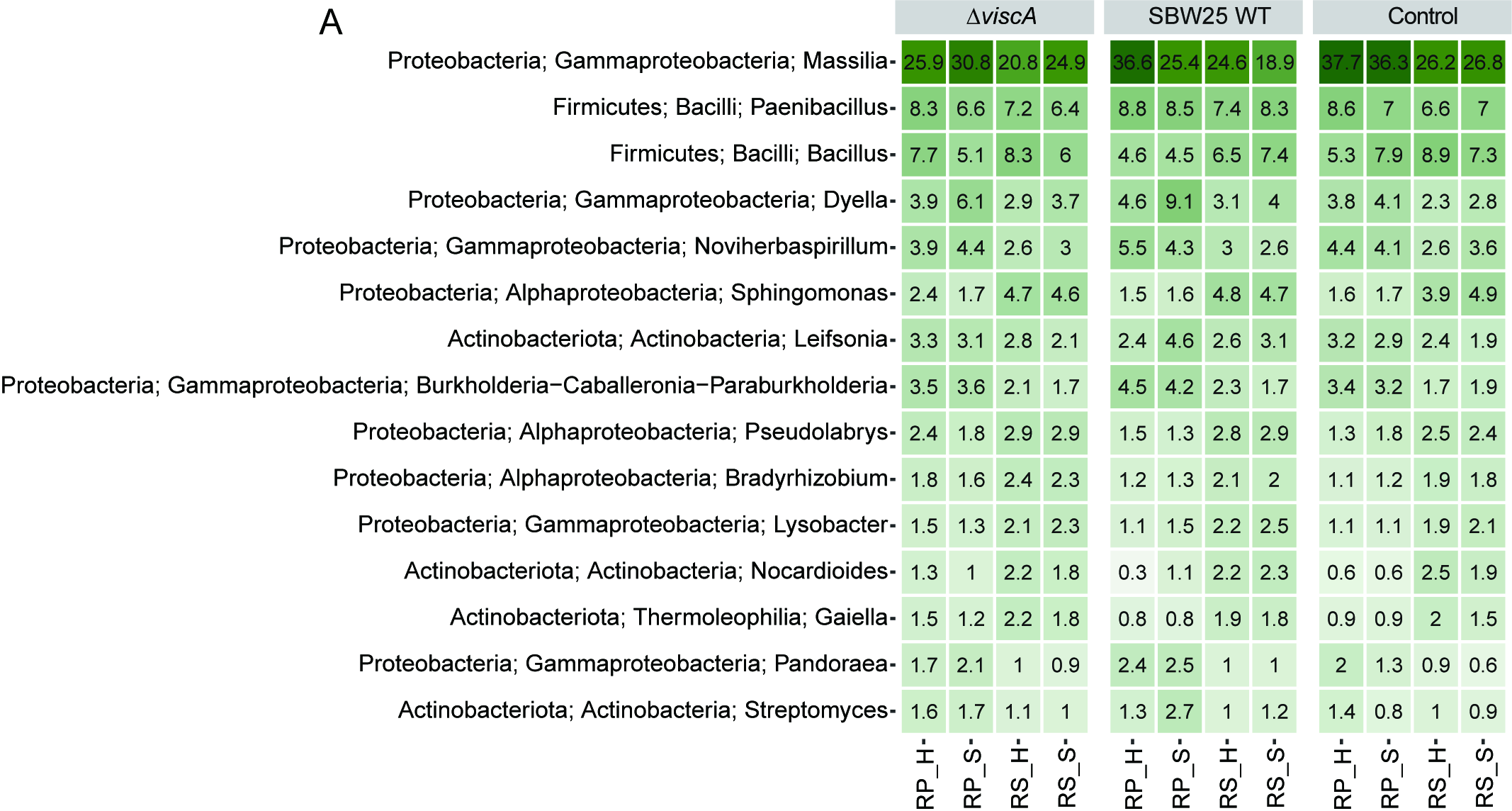

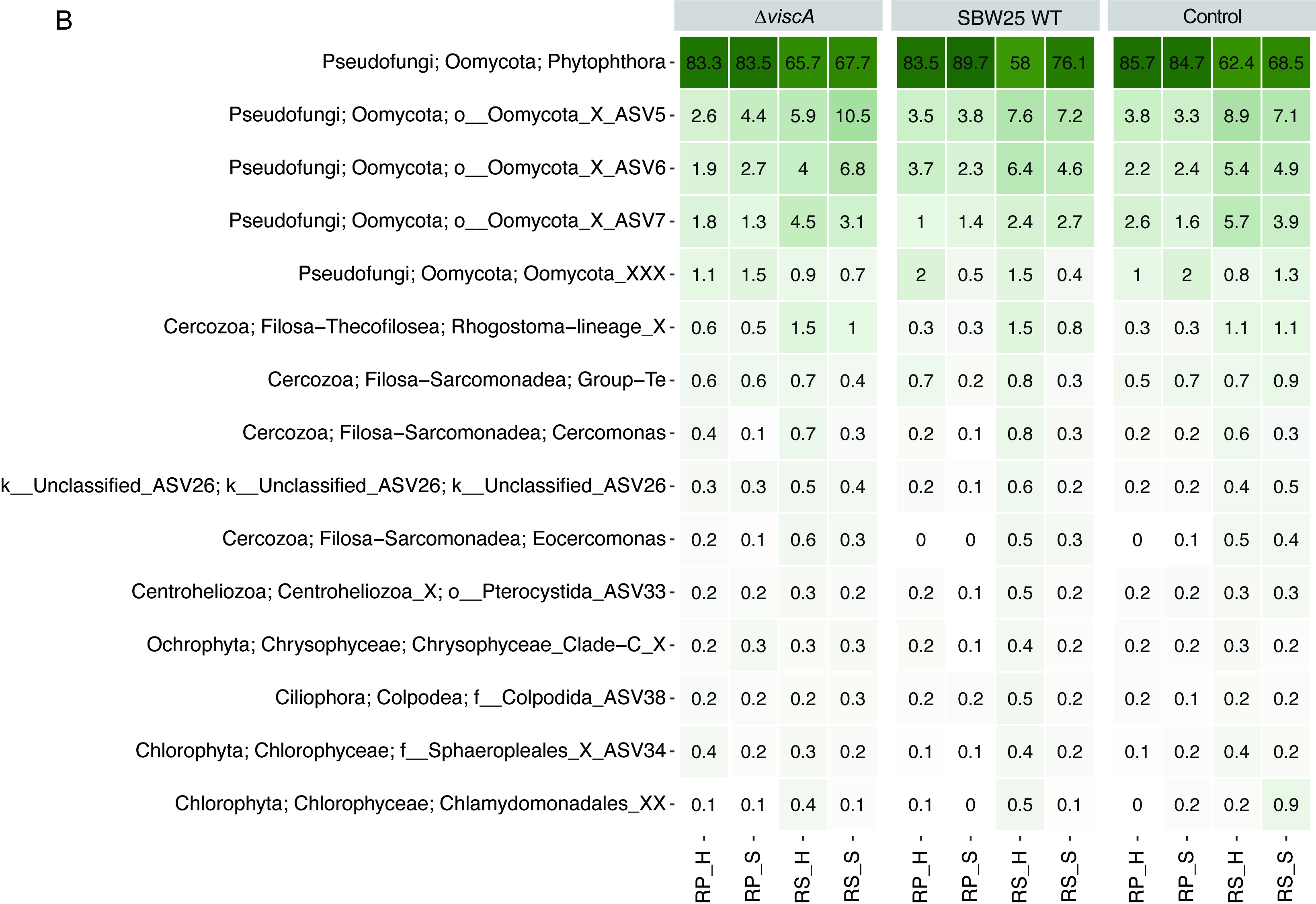
